# Structural insights into the role of Dicer-related helicase 3 in RNAi in *Caenorhabditis elegans*

**DOI:** 10.1101/2021.03.14.433363

**Authors:** Kuohan Li, Jie Zheng, Melissa Wirawan, Nguyen Mai Trinh, Olga Fedorova, Patrick Griffin, Anna Marie Pyle, Dahai Luo

## Abstract

DRH-3 belongs to the family of duplex RNA-dependent ATPases (DRAs), which include Dicer and RIG-I-like receptors (RLRs). DRH-3 is critically involved in germline development and RNAi-facilitated chromosome segregation via the 22G-siRNA pathway in C. elegans. The molecular understanding of DRH-3 and its function in endogenous RNAi pathways remains elusive. In this study, we solved the crystal structures of the DRH-3 N-terminal domain (NTD) and the C-terminal domains (CTDs) in complex with 5’-triphosphorylated RNAs. The NTD of DRH-3 adopts a distinct fold of tandem Caspase Activation and Recruitment Domains (CARDs) structurally similar to the CARDs of RIG-I and MDA5, suggesting a signaling function in the endogenous RNAi biogenesis. The CTD preferentially recognizes 5’-triphosphorylated double-stranded RNAs bearing the typical features of secondary siRNA transcripts. The full-length DRH-3 displays unique structural dynamics upon binding to RNA duplexes that differ from RIG-I or MDA5. These unique molecular features of DRH-3 help explain its function in RNAi in worms and the evolutionary divergence of the Dicer-like helicases.

## INTRODUCTION

RNA interference (RNAi) is important for the viability and growth of *Caenorhabditis elegans* (*C. elegans*) larvae (1–4). RNAi refers to sequence-specific gene silencing that is triggered by small duplex RNAs. These noncoding RNA molecules are key players in RNAi. The two pathways of generating small RNAs in *C. elegans* are cleavage of double-stranded RNA (dsRNA) by Dicer and *de novo* synthesis of secondary siRNAs by worm RNA-dependent RNA polymerase (RdRp). Worm Dicer (DCR-1) is required to produce microRNAs (miRNAs) and small interfering RNAs (siRNAs), which are short (21–23 nt) dsRNAs bearing 5-phosphate (5’-p) ends and 3’ overhangs (5–7). Worm-specific RdRps can transcribe endogenous siRNAs using mRNA templates (2,8,9). These secondary siRNAs exhibit a distinct 5’ to 3’ polarity and bear triphosphate moieties at the 5’ end (5’-ppp). In worms, secondary siRNAs amplify the signal for targeting expressed sequences, which makes them more effective than primary siRNAs in RNAi (10). DRH-3 was identified as a cofactor of worm-specific RdRps and is required for the biogenesis of secondary siRNAs, including endogenous 22G and 26G siRNAs and exogenous secondary virus-derived small interfering RNAs (vsiRNAs) (2,3,11,12).

DRH-3 displays a domain architecture that is similar to other RLRs and includes an N-terminal domain (NTD), a DExH/D helicase core, and a C-terminal domain (CTD) (13–16). The NTD of DRH-3 has no defined function in worm RNAi, making it particularly important to obtain structural information on this enigmatic domain. The NTD shares low amino acid sequence identity with the N-terminal tandem caspase activation and recruitment domains (CARDs) of retinoic acid-inducible gene I (RIG-I) (11%) and melanoma differentiation-associated gene 5 (MDA5) (12%) (14,15). The helicase domain of DRH-3 contains specialized motifs and sequences, indicating that it is related to Dicer and RIG-I-like receptors (RLRs) [namely, RIG-I, MDA5 and laboratory of genetics and physiology 2 (LGP2)](17–20)

Together, these motor proteins are referred to as duplex RNA-dependent ATPases (DRAs) because they bind to dsRNA and hydrolyze ATP but do not unwind dsRNA, which is inconsistent with the canonical function of helicases that separate double-stranded RNA (dsRNA) in the presence of ATP(18). Interestingly, a recent study suggested evolutionary connections between TRIM (tripartite motif) proteins and RNA helicases such as RIG-I and Dicer (21).

The CTD of DRH-3 is homologous to the CTD of RLRs (22,23). Despite this apparent conservation in the overall CTD structure, the RNA recognition specificity is different within this family of proteins, which has important functional implications for viral RNA sensing by RLRs (24–28). For example, the RIG-I CTD shows high binding affinity to the 5’-triphosphate/diphosphate-(5’-ppp/5’-pp) terminus of short dsRNA species (23–25,29–32), whereas the MDA5 CTD preferentially recognizes the internal stem region of long dsRNA and interacts with weaker affinity (27,33,34). The LGP2 CTD can bind both the stem and termini of dsRNA (26). Similar to these related proteins, the DRH-3 CTD may recognize specific RNA features in a way that contributes to its function in the endogenous RNAi pathway.

Structural and functional studies of Dicer proteins and RLRs have greatly enhanced our understanding of their roles in RNAi and antiviral defense mechanisms. In contrast, the molecular mechanisms of endogenous RNAi pathways and the role played by DRH-3 are not well understood. Here, we report crystal structures of the DRH-3 NTD and the CTD in complex with 5’-triphosphorylated RNA. In addition, we monitored the overall conformation and dynamic behavior of the full-length DRH-3 protein in solution using hydrogen/deuterium exchange (HDX) mass spectrometry (MS). Together, our data provide new insights into the specificity and behavior of DRH-3, which will guide functional analyses of the endogenous RNAi pathway in worms.

## MATERIAL AND METHODS

### Protein expression and purification

The cDNA encoding the full-length *C. elegans* DRH-3 (UniProt Q93413) was inserted into the bacterial expression vector Champion pET-SUMO (Invitrogen). The resulting plasmid encodes full-length DRH-3 (DRH-3FL) with an N-terminal hexahistidine (His6)-SUMO tag. The full-length DRH-3 gene was also used as a template for generating recombinant constructs of individual domains of DRH-3. Constructs were designed based on a bioinformatics analysis and secondary structure predictions carried out using BLAST (35), GlobPlot (36) and JPred (37). The His6-SUMO-tagged N-terminal domain (NTD) of DRH-3 (residues 1-335) was cloned into the expression vector pNIC28-Bsa4 (38). The C-terminal domain (CTD) of DRH-3 (residues 940-1108) was cloned into the expression vector pET15b (Novagen), and the resulting clone carries an N-terminal His6 tag and a thrombin cleavage site. All the cloned DNA sequences were confirmed by DNA sequencing. The recombinant constructs were transfected into chemically competent *E. coli* cells [BL21(DE3) Rosetta II (Novagen), BL21(DE3) Rosetta T1R (Novagen) and BL21-CodonPlus (DE3)-RIL cells (Agilent Technologies)] for protein expression.

The transformed cells were grown in Luria broth (LB) with 50 μg ml^−1^ kanamycin and 35 μg ml^−1^ chloramphenicol at 37°C, and protein expression was induced with a final concentration of 500 mM isopropyl β-D-1-thiogalactopyranoside (IPTG) once the OD600 reached 0.6-0.8. The temperature was lowered to 16°C, and the bacterial cell culture was incubated for an additional 20 h for protein expression. The cell pellet was harvested by centrifugation at 4000×*g* for 15 min at 4°C. The cells were resuspended in a buffer containing 25 mM HEPES pH 7.5, 500 mM NaCl, 10% glycerol and 5 mM β-mercaptoethanol and lysed by a Panda Plus 2000 homogenizer (GEA Niro Soavi). The lysate was clarified by high-speed centrifugation at 20 000×*g* at 4°C for 40 min. The supernatant was then incubated with Ni-NTA resin (BioRad) that had been equilibrated with lysis buffer for 1 h at 4°C. The resin was washed with lysis buffer containing up to 20 mM imidazole, and the bound proteins were eluted with lysis buffer containing 200 mM imidazole. Additional heparin chromatography was used during purification of full-length DRH-3 and DRH-3 CTD to remove nucleic acid contamination. The protein was bound to a 5 ml HiTrap HP column (GE Healthcare), and a linear gradient of increasing NaCl concentration was used to elute the protein fractions. The target protein normally eluted at approximately 400 mM to 500 mM NaCl. The eluted protein was then dialyzed overnight with SUMO protease or thrombin to remove the tag. The cleaved DRH-3FL was then further purified using a Superdex 200 HiLoad 16/600 size-exclusion column (GE Healthcare), and the cleaved DRH-3 CTD and DRH-3 NTD were purified using a Superdex 75 HiLoad 16/600 size-exclusion column (GE Healthcare). Peak fractions were pooled and concentrated to 5-15 mg ml^−1^ in a buffer containing 20 mM HEPES pH 7.5, 150 mM NaCl, 5% glycerol and 2 mM DTT. The protein concentration was determined by absorbance spectroscopy at 280 nm, and the purity was assessed by SDS-PAGE.

The N-terminal His6-SUMO-tagged human RIG-I CTD (residues 802-925) was cloned into the expression vector pNIC28-Bsa4 (38)and was expressed and purified in the same way as the DRH-3 CTD.

### Crystallization, data collection and structure determination

All crystals were grown by the hanging-drop vapor diffusion method. The native DRH-3 NTD crystals were grown at 0.2 M ammonium fluoride (pH 6.2), 20% *w/v* polyethylene glycol 3350, and 5 mM DTT at 4°C. The selenomethionine (SeMet)-incorporated DRH-3 NTD crystals were grown in 0.2 M potassium sodium tartrate tetrahydrate (pH 7.4), 20% w/v polyethylene glycol 3350, and 5 mM DTT at 4°C. The DRH-3 CTD was mixed with single-strand 5’-ppp 8-nt RNA at a 1:1 ratio. Crystals were grown from 0.2 M ammonium acetate, 0.1 M Tris pH 8.5, 25% w/v polyethylene glycol 3350 at 18°C. The DRH-3 CTD was mixed with self-complementary 5’-ppp 12-mer GC-rich RNA at a 2:1 molar ratio. Crystals were grown from 0.1 M Tris pH 7.5, 30% w/v polyethylene glycol 6000 at 18°C. Crystals of CTD-RNA complexes were flash-frozen with the reservoir solution and 25% (v/v) glycerol as cryoprotectant. Diffraction intensities for native DRH-3 NTD and SeMet DRH-3 NTD were collected at SLS-PXIII using the PILATUS 2 M-F detector and the multiaxis PIRGo goniometer (Paul Scherrer Institute, Villigen, Switzerland). Integration, scaling and merging of intensities were carried out using the programs iMOSFLM (39) and SCALA (40) from the CCP4 suite (41). Diffraction intensities for CTD-RNA complexes were recorded at NE-CAT beamline ID-24 at the Advanced Photon Source (Argonne National Laboratory, Argonne, IL, USA). Integration, scaling, and merging of the intensities were carried out with the programs XDS (42) and SCALA (43). The structure of the DRH-3 NTD was determined using the single-wavelength anomalous dispersion (SAD) method with the online server AutoRickshaw (44,45). The structure of the DRH-3 CTD in complex with the single-strand 5’-ppp 8-nt RNA was solved by the SAD method using SHELX (46). The structure of the DRH-3 CTD in complex with the 5’ppp dsRNA12 was solved by molecular replacement using Phaser (47). The models were further developed using rounds of manual building in COOT (48) and refined using Phenix (49). Statistics on the data collection and structure refinement for all three structures are summarized in **Table 1**.

**Table 1.** Data collection and Refinement statistics.

| Data collection |  |  |  |  |
| --- | --- | --- | --- | --- |
| Macromolecule | DRH-3 NTD (Native) | DRH-3 NTD (SeMet) | DRH-3 CTD: 5'ppp 8-nt ssRNA | [DRH-3 CTD: 5'ppp 12-bp dsRNA]2 |
| Wavelength (Å) | 0.9998 | 0.9794 | 1.2819 | 0.9792 |
| Resolution (Å) | 85.56 - 2.80 (2.95 - 2.80) | 47.63-3.11(3.28-3.11) | 43.04 - 1.89 (1.93 - 1.89) | 43.96 - 1.60 (1.63 - 1.60) |
| Space group | P 41 21 2 | P 41 21 2 | P 43 21 2 | P 2 21 21 |
| Unit cell a/b/c(Å) | 93.18/93.18/215.96 | 89.93/89.93/215.68 | 56.843/56.843/131.75 | 33.775/93.090/133.823 |
| $\alpha/\beta/\gamma$ (°) | 90/90/90 | 90/90/90 | 90/90/90 | 90/90/90 |
| Unique reflections | 24012 (3401) | 16670(2324) | 17289 (984) | 56358 (2845) |
| Multiplicity | 11.1 | 12.8 | 9.2 | 3.1 |
| Completeness (%) | 99.3 (99.0) | 99.7(98.3) | 96.3 (88.4) | 99.4 (99.9) |
| I/sigma(I) | 10.9 (2.2) | 11.0(2.1) | 12.0 (1.7) | 14.9(1.6) |
| R<sub>merge</sub> | 0.134 (0.896) | 0.241(1.352) | 0.142 (1.715) | 0.042 (0.761) |
| Wilson B-factor | 66.9 | 59.3 | 26.7 | 20.1 |
| CC1/2 | 1 (0.727) | 0.996(0.664) | 0.996 (0.378) | 0.998 (0.586) |
| Refinement Statistics(\|F\|>0) |  |  |  |  |
| Reflections used in refinement | 22582 |  | 17263 | 56281 |
| Number of atoms | 5390 |  | 1678 | 3650 |
| macromolecules | 5252 |  | 1521 | 3188 |
| ligands | 0 |  | 33 | 130 |
| water | 138 |  | 124 | 332 |
| Protein residues | 659 |  | 167 | 322 |
| R<sub>work</sub> | 0.2337 |  | 0.1777 | 0.1692 |
| R<sub>free</sub> | 0.286 |  | 0.2213 | 0.1988 |
| Rmsd from ideal bond length(Å) | 0.008 |  | 0.008 | 0.017 |
| Rmsd from ideal bond angle(°) | 1.19 |  | 0.991 | 1.88 |
| Ramachandran statistics |  |  |  |  |
| Ramachandran favored (%) | 97.6 |  | 95.73 | 98.2 |
| Ramachandran allowed (%) | 100 |  | 100 | 100 |
| Ramachandran outliers (%) | 0 |  | 0 | 0 |
| Clashscore | 6.53 |  | 2.66 | 5.53 |
| Average B-factor | 86.3 |  | 37.06 | 29.77 |
| macromolecules | 86.8 |  | 36.32 | 28.96 |
| solvent | 67.43 |  | 38.98 | 36.60 |

a. Values in parentheses indicated values in the highest resolution shell b. $R_{\text{merge}} = \Sigma |I_{\text{obs}} - I_{\text{avg}}| / \Sigma I_{\text{avg}}$ . c. $R_{\text{free}} = \Sigma |F_{\text{obs}} - F_{\text{calc}}| / \Sigma I_{\text{calc}}$ . $R_{\text{free}}$ was calculated with 5% of reflections.

### RNA preparations

The 5’-ppp RNAs used for crystallization and HDX/MS were synthesized using T7 polymerase *in vitro* transcription. Transcription was carried out using template DNA with a T7 promoter (5’-G TAATACGACTCACTATA…), and the sequence was initiated by GTP. The reaction was then phenol-chloroform extracted, precipitated with 80% ethanol and purified on a 20% denaturing gel. The transcribed RNAs were run on a 20% denaturing gel to test for purity and size. All palindromic dsRNA oligonucleotides used for assays were synthesized on a MerMade 12 DNA-RNA synthesizer (BioAutomation) as previously described (50,51). Synthesized RNAs were deprotected (50) and gel purified. The synthetic RNAs were further analyzed for purity by mass spectrometry (Novatia LLC). FAM-labeled RNA was purchased from Integrated DNA Technologies. All RNA sequences are listed in Supplementary **Table S2**.

### Limited Proteolysis

A 1 mg ml^−1^ trypsin stock solution was prepared in trypsin digestion reaction buffer (25 mM HEPES, 5% glycerol, 150 mM NaCl, and 1 mM DTT). DRH-3 CTD proteins and DRH-3 CTD-RNA complexes were incubated with trypsin at 1:2000 w/w for 0-2 hours at 25°C. The reactions were quenched by the addition of SDS loading dye and boiling for 5 min. Samples were analyzed by SDS-PAGE.

### Thermal shift assay

The thermal shift assay involves monitoring changes in the fluorescence signal of SYPRO Orange dye as it interacts with a protein undergoing thermal unfolding. The SYPRO Orange dye was supplied by Invitrogen (catalog number S6650) at 5000x concentration in DMSO and diluted in assay buffer for use. Each test sample (50 μl total volume) in a 96-well PCR plate (BioRad) contained 25 mM HEPES (pH 7.4), 150 mM NaCl, 1 mM DTT, 5% glycerol, 5x SYPRO Orange dye, 5 μM enzyme, and 6 μM RNA. In negative control samples, buffer was added instead of test RNA(s). The plate was sealed with optical quality sealing tape (BioRad) and heated in an i-Cycler iQ5 real-time PCR detection system (BioRad) from 25 to 95°C in increments of 1°C. Fluorescence changes in the wells were monitored simultaneously with a charge-coupled (CCD) camera. The wavelengths for excitation and emission were 485 and 575 nm, respectively. The temperature midpoint for the protein unfolding transition, the melting temperature *(Tm),* was calculated using BioRad iQ5 software.

### Fluorescence polarization (FP) competition assay

Fluorescence polarization experiments were performed on a Synergy H1 plate reader (BioTek, Vermont) in 25 mM HEPES, 2.5% glycerol, 75 mM NaCl, 0.5 mM DTT, and 0.01% Triton-X. The excitation and emission wavelengths were set to 485 and 510 nm, respectively. The 5’-OH hairpin RNA was purchased from IDT, and fluorescein amidite (FAM) was labeled at the hairpin region of the RNA. For competition assays, RNA competitors were serially diluted over a concentration range from 10 μM to 2.4 nM. DRH-3 CTD (400 nM), FAM-labeled RNA (5 nM) and RNA competitors were mixed and distributed equally in a black, 384-well, low-flange microplate (Corning, UK). The inhibition (%) was calculated using the equation below.

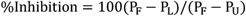

where P_F_ is the polarity of fully bound 5’-OH hairpin RNA;

P_U_ is the polarity of unbound 5’-OH hairpin RNA; and

P_L_ is the polarity of 5’-OH hairpin RNA in the presence of RNA competitors.

### Hydrogen/deuterium exchange coupled with mass spectrometry (HDX-MS)

DRH-3FL proteins were exposed to deuterated water for 1 hour followed by a quenching step (low pH on ice) to stop hydrogen/deuterium exchange. Proteins were digested into peptide fragments by low-temperature and low-pH compatible pepsin. The resulting peptides were subsequently separated by capillary-flow liquid chromatography and measured by mass spectrometry. The level of deuterium incorporation was then determined by examining the change in the mass centroid. HDX Workbench (52) was utilized to analyze the data and generate graphs. HDX-MS results for DRH-3 proteins shown as a graphical heatmap that revealed the level of deuterium incorporation (%D) for detected peptides. The heatmap of HDX was subsequently projected onto the crystal structure of DRH-3 CARDs and DRH-3FL models with and without RNA ligand to represent the molecular structure and protein dynamics of DRH-3.

### Structure modeling

Homology models of DRH-3 proteins were generated using SWISS-MODEL (53). DRH-3 CARD (PDB code: 6JPX), RIG-I helicase domain (PDB code: 5E3H) and DRH-3 CTD (PDB code: 6JPY) crystal structures were used as input reference models. For illustration purposes, the HDX-MS data were taken into account to model the relative positions of the domains.

### Negative staining transmission electron microscopy

The samples were prepared as follows: 50 μg/mL of DRH-3 FL recombinant protein were mixed with 3 μg/mL of dsRNA of different length in buffer containing 25 mM HEPES pH 7.5, 0.1 M KCl, 5 mM MgCl2, 1.5 mM ADP-AlF4. RNAs used in this experiment are listed in Supplementary **Table S2**.The mixture were then incubated on ice for 30 min. After incubation, 4μL of each sample were applied to glow-discharged carbon-coated grids. The grids were then negatively stained with uranyl acetate [1% (wt/vol)] and imaged with an FEI Tecnai 12 transmission electron microscope at an accelerating voltage of 200 kV. Micrographs were taken at −1 μm defocus, 49000× magnification and 3.061Å per pixel.

## RESULTS

### DRH-3 NTD contains unique tandem CARDs in a parallel arrangement

The DRH-3 NTD crystal structure was solved at 2.8 Å resolution (**Figure 1**), revealing a domain that is composed primarily of α helices arranged in two tandem helical bundles (**Figure 1C**). The two bundles adopt Greek-key-like folds linked by loops and small helices (54,55). We compared the structures of the α-helical bundles with their respective top 5 hits from the DALI server (56) (**Figure S1 and Table S1)**. All hits fell into the CARD domain family of proteins that participate either in apoptosis [nucleolar protein 3 (NOL3), procaspase-9 (pc-9) and apoptotic protease activating factor 1(Apaf-1)] or inflammation [caspase recruitment domain family member 11(CARD11), ICEBERG, RIG-I, B Cell CLL/Lymphoma 10 (Bcl10) and Bcl10-interacting CARD protein (BinCARD)]. Despite the low sequence identities (less than 10%) shared by the two α-helical bundles of the top hits, the relative orientation of the helices is similar. We therefore refer to the two helical bundles in the DRH-3 NTD as CARD1 (helix α1-6) and CARD2 (helix α7-12). The first helix in DRH-3 CARD1 has a kink, which is a common feature for CARDs (57,58). The third and fourth helices in DRH-3 CARD1 and CARD2 are extremely long and different from canonical CARDs (59). The two CARDs in DRH-3 are sequentially (11% amino acid sequence identity) and structurally (rmsd of 4.9 Å for all Cα atoms) distinct, and they are stabilized side-by-side by hydrogen bonds between helices α1-α10 and helices α4-α7 (**Figure S2)**. In contrast, the two CARD domains of RIG-I are similar to each other (20% amino acid sequence identity), and they are oriented in a head-to-tail arrangement.

**Figure 1.**
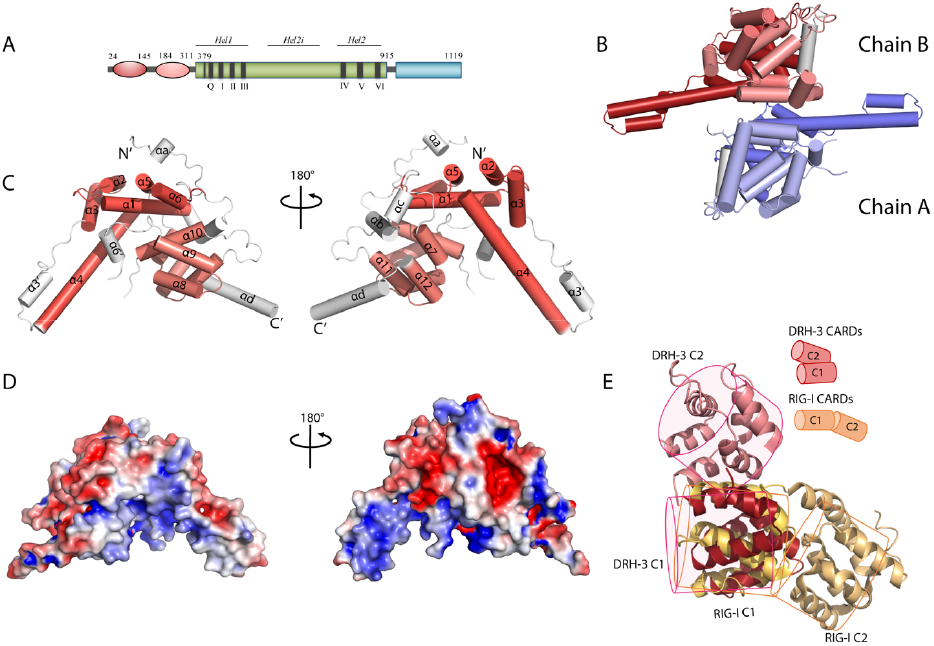
Overall structure of DRH-3 N-terminal domain. (A) Schematic representation of the domain architecture of DRH-3. NTD, worm-specific N-terminal domain; HEL, the helicase domain; CTD, the C-terminal RNA binding domain. The residues numbers corresponding to the predicted domain boundaries were labelled accordingly. (B) Crystal structure of dimeric DRH-3 NTD. Two DRH-3 NTD molecules in the asymmetric unit are shown as cartoons. Chain A and Chain B are indicated by different colours. (C) Cylinder depiction of DRH-3 NTD structure with helices labelled. DRH-3 NTD contains two CRAD like α-helical bundles, highlighted in red (NTD_C1) and pink (NTD_C2) colours, respectively. The topology of a 6-α-helical bundle is that of a Greek Key. Cylinders represent the helices and the arrow represents the rotation direction of the polypeptide chain. (D) Surface of DRH-3 NTD coloured according to the electrostatic charges where blue is for positive charges, red for negative charges, and white for neutral surface. (E) Superposition of DRH-3 NTD1 with RIG-I CARD1. The relative position of the two CARD/CARD-like domain in DRH-3 and RIG-I are compared and illustrated in simplified carton models.

We analyzed the electrostatic potential at the solvent-accessible surface of DRH-3 CARDs (**Figure 1E**). The surface of DRH-3 CARDs contains several highly charged patches. The positive patches fall onto the helix stem composed of helices α4 and α3’; the negative patches were on the upper helix stem region (helices α2 and α3). These surfaces may contribute to the functions of DRH-3 because death-fold proteins signal via electrostatic interactions (58). Two identical DRH-3 NTD molecules (rmsd 0.367 Å for 305 superimposed Cα atoms) were present in the asymmetric unit. To examine whether the dimeric interface was biologically relevant, we analyzed the structure using PDBePISA (60) (**Figure S3**). The interface area (540 Å^2^) is approximately 10% of the total solvent-accessible area (5273 Å^2^), suggesting that dimer formation is likely the result of crystal packing. Indeed, both DRH-3 CARDs and DRH-3FL exist as monomers in solution (**Figure S4**). Overall, the NTD of DRH-3 folds into a set of novel tandem CARDs, which may function by interacting with certain CARD-containing protein(s) in warm RNAi.

### DRH-3 CTD recognizes 5’ tri-phosphorylated dsRNA

To elucidate the structural basis for secondary siRNA recognition, we identified two crystal structures of the DRH-3 CTD in complex with 5’-ppp RNAs (**Table 1**). The overall conformation of the DRH-3 CTD was similar to that of the RLR CTDs: they all contain a saddle-like shape that consists of two antiparallel β-sheets (β-sheet1: β1-2 and β9-10, β-sheet2: β5-8) and exhibits a high degree of shape complementarity to the RNA ligands. The long helix (α4) at the C-terminus, which stacks on the back side of the 5’-ppp binding site, is a unique feature of DRH-3. A C4-type zinc finger binding motif (strands β1-2 and β6-7) is found in the DRH-3 and RLR CTDs, and it captures a zinc ion that plays a role in maintaining the overall fold of the CTD (24–27).

5’-ppp participates in an extensive network of electrostatic interactions with the DRH-3 CTD (**Figure 2 and S5**). For example, K992 and R1016 interact with γ-phosphate; K990 and K993 interact with γ- and β-phosphates; and K988 interacts with α- and β-phosphates. Similar charged residues within the CTD of RIG-I are important for binding 5’-ppp RNAs (K858-K861)(24,25,30). In addition, residues K1048 and S1050 in the conserved KWK motif (61,62) mediate interactions between the DRH-3 CTD and the RNA sugar phosphate backbone. The 3’-stranded RNA contributes to RNA binding via a hydrogen bond between C12 and residues F969. Notably, the DRH-3 CTD binds to 5’-ppp dsRNA and 5’-ppp ssRNA in a similar manner (**Figure 2 and S5)**.

**Figure 2.**
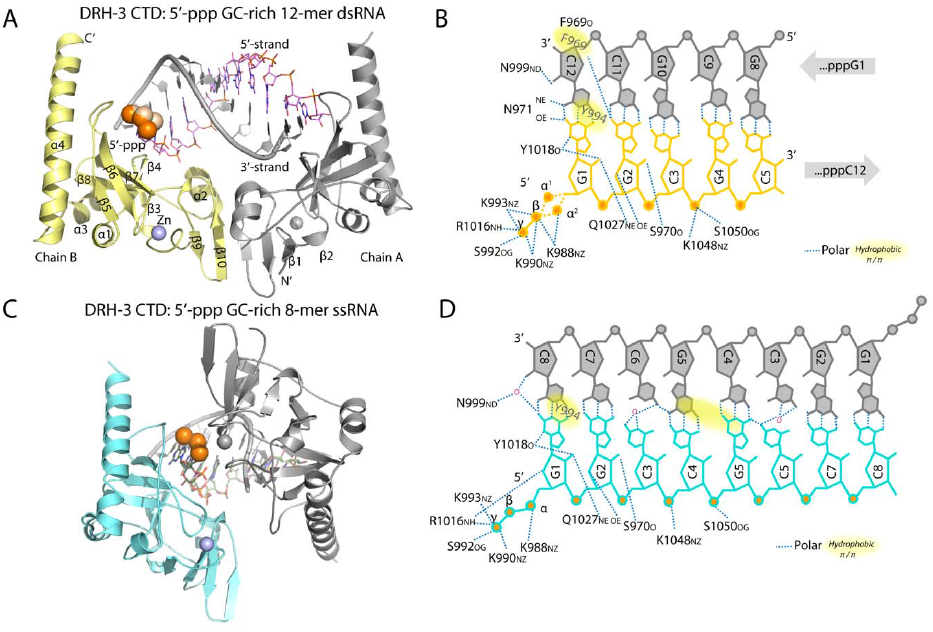
5’-ppp RNA recognition by DRH-3 CTD. (A) Overall structure of DRH-3 in complex with 5’-ppp 12-mer dsRNA. The crystallographic asymmetric unit contains a 2:1 complex between DRH-3 CTD and the dsRNA. Chain B and the 5’ stand are coloured. The 5’-ppp is shown as orange spheres and the zinc molecule is shown as a purple sphere. (B) The schematic diagram of contacts between DRH-3 CTD and 5’-ppp 12-mer dsRNA. (C) Overall structure of DRH-3 in complex with 5’-ppp 8-mer ssRNA. The crystallographic asymmetric unit contains a 1:1 complex between DRH-3 CTD and the ssRNA. The symmetry mate structure is shown in grey. The 5’-ppp is shown as orange spheres and the zinc molecule is shown as a purple sphere. (D) The schematic diagram of contacts between DRH-3 CTD and 5’-ppp 8-mer ssRNA.

To further interpret these observations, the electrostatic potential at the solvent-accessible surface of DRH-3 CTD was analyzed. A continuous positively charged surface area was identified, and it corresponds to the 5’-ppp recognition and the RNA phosphate backbone binding sites (**Figure 2**). We then compared the DRH-3 CTD with the available human RLR structures (**Figure 3**). Superposition of the DRH-3 CTD with the RLR CTDs resulted in a rmsd value ranging from 0.9 to 2.3 Å for all superimposed Cα atoms, indicating a high level of structural similarity despite a relatively low sequence similarity. Intriguingly, the 5’-ppp binding loop between β-strands 5 and 6 is similar in DRH-3, RIG-I and LGP2, which is consistent with their preferences for polyphosphorylated dsRNA 5’ termini.

**Figure 3.**
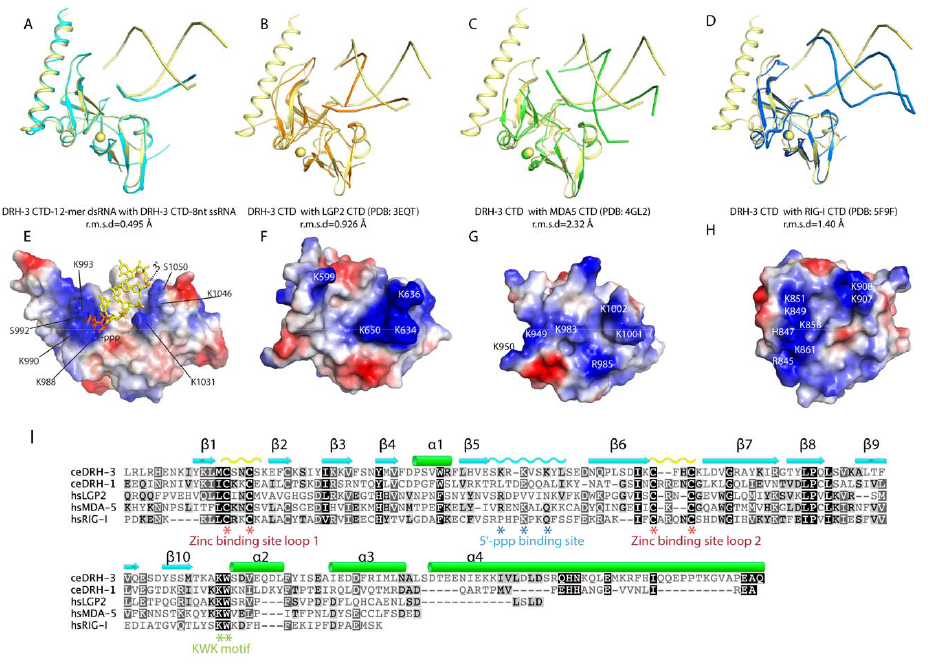
Comparison of C-terminal domain of DRH3 and RLRs. (A) Superposition of DRH-3 CTD structures bound to the 5’-ppp 12-mer dsRNA and the 5’-ppp 8-mer ssRNA. (B) Superposition of the RNA bound DRH-3 CTD (yellow) and LGP2 CTD (orange, PDB code: 3EQT). (C) Superposition of the RNA bound DRH-3 CTD (yellow) and MDA5 CTD (green, PDB code: 4GL2). (D) Superposition of the RNA bound DRH-3 CTD (yellow) and RIG-I CTD (orange, PDB code: 3EQT).Polypeptide chains are shown as cartoons. Structural superimposition was done by pairwise and represented by Pymol [43]. (E) Electrostatic surfaces of dsRNA-bound DRH-3 CTD. Positively charged surfaces are coloured blue and negatively charged surfaces are red. Key residues involved in the RNA binding were labelled. (F) Electrostatic surfaces of LGP2 CTD. (G) Electrostatic surfaces of MDA5 CTD. (H) Electrostatic surfaces of RIG-I CTD. Each CTD is shown in the same orientation as in Figure 3 A-C. Positively charged surfaces are coloured in blue and negatively charged surfaces are coloured in red. The key charged residues are indicated. (I) Alignment of amino acid sequences of DRH and RLR CTDs. Sequences were aligned using the program ClustalW and visualized using Geneious software. The conservation of the amino acids is indicated by colours.

To determine the RNA ligand binding characteristics of DRH-3, we tested whether the 5’-modifications (5’-ppp, 5’-OH and 5’-p) and 5’ or 3’ overhanging regions on the 5’-ppp dsRNA affect RNA binding to the DRH-3 CTD (**Table S2**). First, we examined whether the bound RNA can protect the DRH-3 CTD from trypsin digestion **(Figure 5A)**. Unbound DRH-3 CTD was digested into two major fragments after 5 minutes of treatment with trypsin. When the DRH-3 CTD was incubated with different dsRNA species prior to treatment with trypsin for 5 minutes, the digestion patterns were very similar, suggesting that the RNA did not protect the DRH-3 CTD from trypsin digestion. In contrast, all the 12-mer dsRNAs protected RIG-I CTD from trypsin digestion, especially blunt-ended 5’-ppp and 5’-pp and 5’-ppp with a 3’-overhang, suggesting that RIG-I CTD prefers to bind to 5’-ppp RNAs **(Figure 5B)**. We also used a thermal stability assay to study the effect of RNA binding on protein stability. No significant difference in the melting temperature (*T_m_*) was observed in the presence and absence of the RNA ligands, indicating that the binding of RNAs failed to change the thermal stability of the DRH-3 CTD. In contrast, all RNAs, especially the 5’-ppp with a 3’-overhang, increased the thermal stability of the RIG-I CTD, as expected (**Figure 5D**). Together, the limited proteolysis and thermal stability methods both suggested that the DRH-3 CTD does not display a strong specificity for a particular type of RNA 5’-terminus. To evaluate this aspect more carefully, we monitored RNA binding by fluorescence polarization (FP) using a competition assay. The binding affinity of the DRH-3 CTD to the RNA probe, which is a FAM-labeled 5’-OH hairpin RNA, was determined to be 325 nM (Fig S9). We then titrated each competitor RNA through a range of concentrations that resulted in a maximum polarity of 50% (IC50 values) (**Figure 5E**). Consistent with the crystallographic findings that showed apparent interactions between 5’-ppp RNA ligands and specific amino acids of the DRH-3 CTD, 5’-ppp RNAs showed the highest degree of competition with the probe (IC50 values ranging from 80 nM to 234 nM), indicating that the DRH-3 CTD preferably recognized 5’-ppp RNAs. In contrast, the 5’-OH and 5’-p RNAs competed poorly with the RNA probe, showing relatively high IC50 values (631 nM to 752 nM). Overall, the CTD of DRH-3 apparently preferentially recognizes the 5’ppp blunt-ended dsRNAs, which is consistent with the conclusions from previous studies using DRH-3FL proteins (13,28)

### DRH-3FL displays an open conformation and binds to 5’ppp dsRNA via HEL-CTD but not CARDs

To examine the overall architecture of DRH-3FL in response to different ligands, we performed hydrogen/deuterium exchange coupled with mass spectrometry (HDX-MS). HDX-MS was previously used to dissect the overall conformational changes of RIG-I and MDA5 (63–65). The exchange rates of amide hydrogens in the target protein in the presence of deuterated solvent were mapped to the primary amino acid sequence and structural models of DRH-3 conformational states. First, the CARDs within the NTD in isolation and CARDs of the DRH-3FL free protein showed similar deuteration profiles, indicating that there are no intramolecular interactions between the CARDs and the core central domains of DRH-3FL. This finding is different from what is observed with RIG-I, in which the CARDs are sequestered in an autoinhibitory conformation (17,64,66) (**Figure 4A-B and S6-7**). The majority of peptides on DRH-3 CARDs (86%) exhibited a magnitude of deuterium uptake that was less than 50%. This low level of H/D exchange suggests a rigid conformation of the DRH-3 CARDs. In comparison, the linker region (I351-E368) between the CARDs and the helicase domain (HEL) was more solvently exposed (70% deuteriation) in DRH-3FL. Therefore, the overall conformation of DRH-3FL appears to be similar to that of free MDA5, in which the various protein domains are relatively unconstrained in the free form (64).

**Figure 4.**
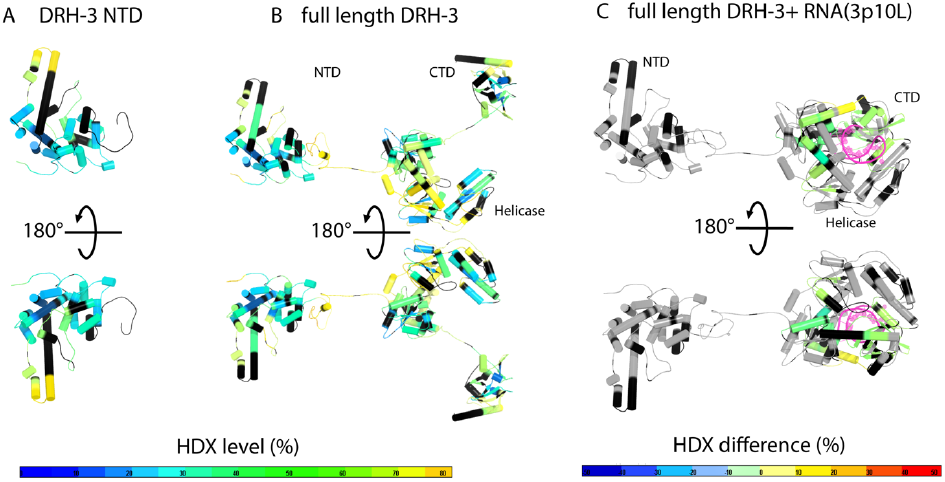
Conformational dynamics of DRH-3. Deuterium uptake profiles of DRH-3 NTD (A) and full length DRH-3 (B) are presented in a front and back view. The structure of DRH-3 NTD was obtained from this study and the full length DRH-3 model was generated by assembling all three domains via fusing peptide bonds. The deuterium exchange data was mapped onto the structural models of DRH-3 NTD and full length DRH-3. The % deuterium exchange in the deuterium exchange of each peptide is coloured according to the scale bar. Blue indicates the lowest and orange indicates the highest exchange rate. Black indicates that the region has no amide hydrogen exchange activities. (C) HDX-MS differential map for full length DRH-3 with a short hairpin RNA in a front and back view. Differential single amino acid consolidation HDX data was mapped onto full length RNA-bound DRH-3 structural model. The percentages of deuterium differences are color-coded according to the scale bar.

**Figure 5.**
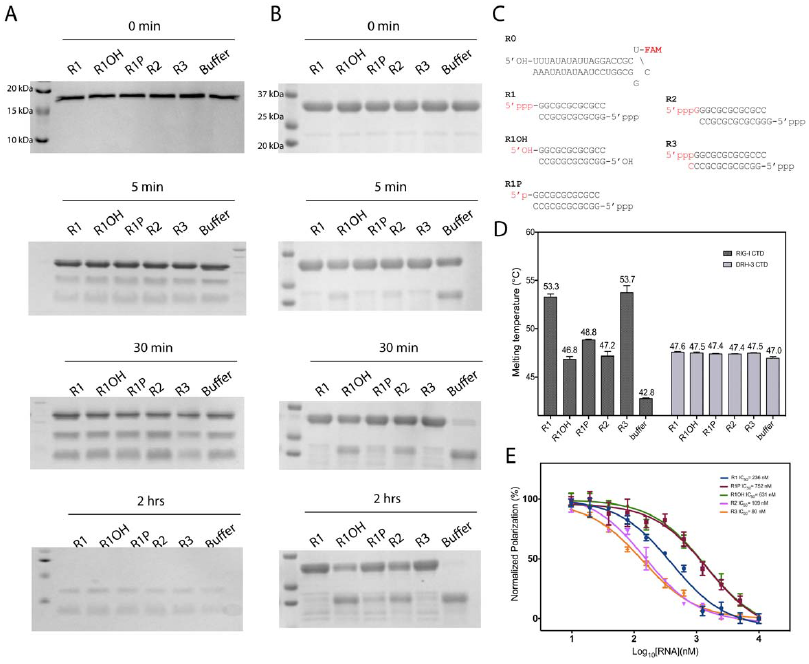
RNA ligand binding properties of DRH-3 CTD. (A-B) Proteolysis effect of RNAs on CTDs. DRH-3 CTD (A) and RIG-I CTD (B) were digested with trypsin at different time periods, and quenched by adding SDS protein loading dye. Denatured proteins were resolved on reducing SDS-PAGE. (C) RNA oligonucleotides with different 5’ or 3’ end modifications. (D) Stabilisation of CTDs by RNA binding. Thermal shift assay results for DRH-3 CTD and RIG-I CTD in the absence (buffer as negative control) and the presence of test RNAs. The melting temperature *Tm* values (°C) are shown above the bars. All data are mean ± SEM for two independent experiments done in triplicate. (E) Validation of the RNA binding affinity of DRH-3 by FP competition assay. Percentage inhibition of the FAM-labelled 5’-OH 18-mer hairpin RNA binding versus RNA competitors concentration response curves. Experiments were done in triplicate and average FP value was plotted with one standard deviation of the mean shown as error. The titration curves were fitted using GraphPad prism and IC5os were calculated from the titration curves.

To map the RNA binding interface and the RNA-induced conformational dynamics of DRH-3FL in solution, we then performed HDX-MS experiments for DRH-3FL in the presence of the short 5’-ppp hairpin RNA 3p10L (5’-ppp GGACGUACGUU<u>UCGA</u>CGUACGUCC, also known as SLR10 (67)), which was shown to bind and activate RIG-I to turn on the type I interferon-mediated immune response (32,64,67). In the presence of 3p10L, several conserved motifs within the helicase domain (motifs Ia, II, IVa, Vb and VI) and the CTD (the 5’-ppp binding site and the KWK motif) displayed a significant reduction in deuteration, suggesting their involvement in RNA binding. A helix of Hel2i and the Pincer region were also affected by 3p10L binding (**Figure 4C and S8**). Importantly, the H/D exchange rate of the CARDs in the DRH-3FL-3p10L complex remained unchanged, suggesting that the binding of 3p10L had no allosteric influence on the dynamics of the CARDs, unlike the RIG-I CARDs.

To determine whether DRH-3FL can oligomerize upon long RNA binding. The recombinant DRH-3FL protein were incubated with dsRNA of various lengths (**Table S2**). The samples were then applied on carbon-coated grids and negatively stained with uranyl acetate. As shown in Figure 6, no oligomeric form of DRH-3 was identified in the presence of dsRNA of 100 bp in the micrographs obtained (**Figure 6**). This is unlike MDA5 which forms RNA-associated helical oligomers and cooperative filaments (34,68–71).

**Figure 6.**
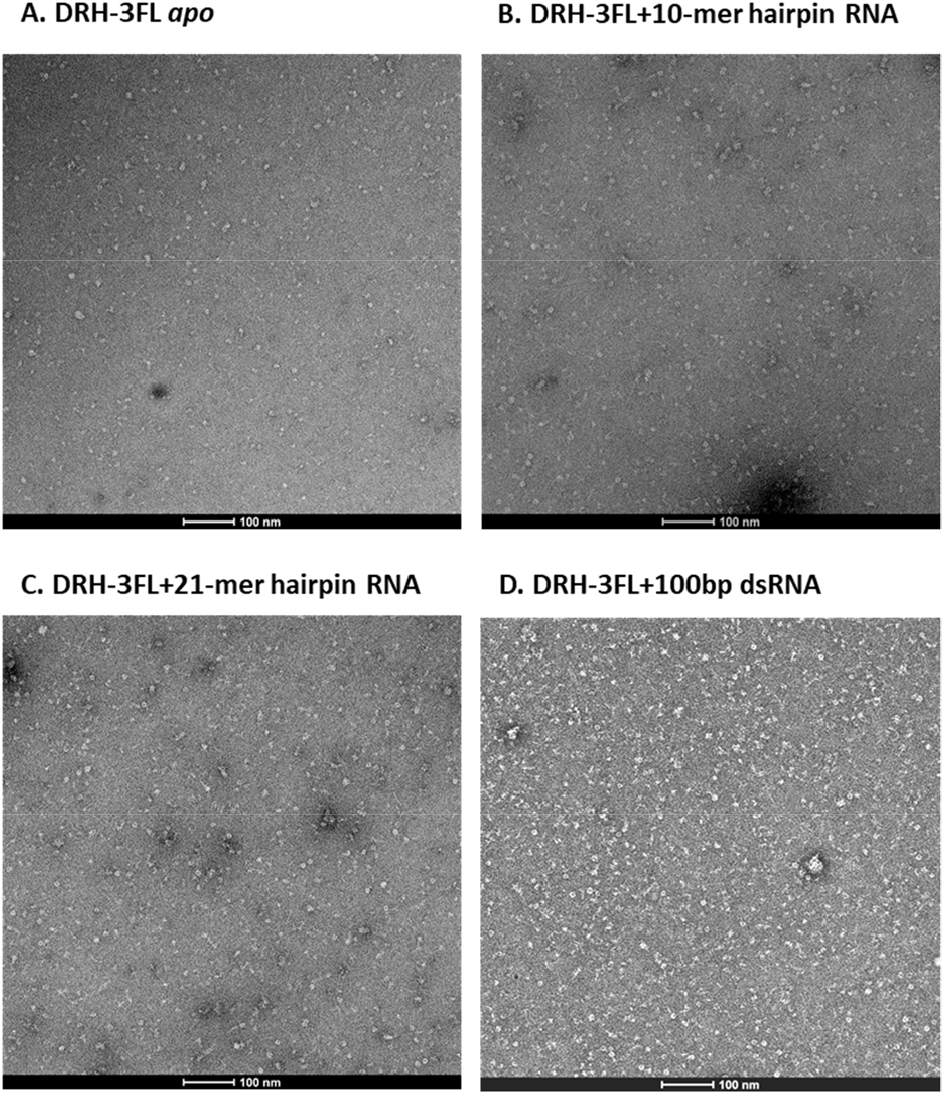
Negative stain electron microscopy images of DRH-3FL dsRNA complexes. (A) Negative stain electron microscopy of *apo* DRH-3FL. Scale bar represents 100 nm. (B-D) Negative stain electron microscopy of RNA-associated DRH-3FL. The recombinant DRH-3FL protein were incubated with dsRNA of 10, 21, and 100bp.

In summary, free DRH-3 adopts an extended conformation where the N-terminal CARDs HEL and CTD are connected by flexible linkers and are individually solvent exposed. DRH-3FL seems to behave differently from both RIG-I and MDA5, as there were no apparent intramolecular autoinhibitory interactions or oligomerization induced upon binding to dsRNA (66,68,71–74).

## DISCUSSION

In this study, we performed structural and functional studies of the DRH-3 protein from *C.elegans.* The crystal structure of the DRH-3 N-terminal domain is observed to be structurally similar to death-fold proteins, especially CARDs. In mammals, CARDs play important roles in apoptosis, immune responses and cytokine processing signaling pathways, while in worms, CARDs are restricted to apoptotic proteins (55). DRH-3 is the first CARD-containing protein involved in the RNAi pathway that has been discovered thus far. The tandem CARDs of DRH-3 are different from the CARDs of RIG-I because the two CARDs of DRH-3 arrange themselves in a parallel orientation instead of the head-to-tail orientation found in RIG-I. In addition, the DRH-3 NTD exists as a monomer in solution, while the RLR CARDs form homotypic oligomers. CARD-CARD interaction is required for CARD-containing proteins to perform their signaling functions (54). Therefore, if DRH-3 CARDs don’t bind to each other, they might interact with other CARD-containing proteins via heterotypical CARD-CARD interactions during the process of RNAi. Alternatively, they may bind to a completely different type of protein interaction partner.Oligomer formation among CARDs primarily depends on interactions between charged surfaces and is relatively independent of amino acid sequences. As such, the DRH-3 CARDs and their prospective binding partners are expected to exhibit complementarity of charged interfaces.

Our structural studies show a highly positively charged binding site on DRH-3 CTD for 5’-ppp. This 5’-ppp binding site is also conserved in RIG-I CTD (**Figure 3**). In humans, the 5’-ppp dsRNA is a viral pathogen-associated molecular pattern (PAMP) that is specifically recognized up by RIG-I, whose activation triggers downstream Type I interferon responses. The binding affinities of 5’-OH dsRNA and 5’-ppp ssRNA are particularly low compared to that of 5’-ppp dsRNA (24,30,31). ATP hydrolysis facilitates the removal of lower-affinity self-RNAs from the RIG-I binding site (23,31,32,75–77). Similarly, DRH-3 CTD can discriminate different types of RNA ends and may contribute to target RNA selection. The FP-based competition assays carried out in this study show that DRH-3 prefers RNAs that are terminated by a 5’-ppp (a signature of RdRp-generated RNAs) in comparison with 5’-p and 5’-OH, Dicer-cleaved RNAs or other exogenous RNAs, which suggests that it plays a specific role in secondary siRNA recognition. In *C. elegans*, RdRps transcribe secondary small regulatory RNAs *de novo* from mRNA-derived templates (2,8,9). Together with a dual Tudor domain protein, DRH-3 is required for the biogenesis of secondary small RNAs, including endogenous 22G and 26G RNAs and exogenous secondary vsiRNAs (2,3,78,79).

We obtained structural information on DRH-3FL using HDX-MS. The exchange rate of amide hydrogens in the target protein with deuterated solvent was mapped to the primary sequence and the structural model of DRH-3. Oligomerization of CARDs is thought to enhance translocation rates and provide a basis for cooperativity in ATPase hydrolysis and/or translocation (17,18,44,68,80). The activation of CARD oligomerization may require specific activation mechanisms. For example, MDA5 displays strong cooperative binding and forms oligomers on long dsRNA (64). The oligomerization of RIG-I CARDs depends on a conformational change induced by RNA binding and ATP hydrolysis (21,81). Given the involvement of DRH-3 in 22G siRNA biogenesis, it is likely to sense the 5’ triphosphorylated dsRNA 22 base pairs along with other partner proteins, such as worm-specific RdRp and Agos (Aoki et al., 2007, Gu et al., 2009, Nakamura et al., 2007, Vasale et al., 2010).

To our knowledge, DRH-3 is the first identified CARD-containing protein with involvement in the RNAi pathway. Structural studies of DRH-3 CTD provide important insights into the basis of secondary siRNA recognition of DRH-3. However, further studies are needed to fully characterize the role of DRH-3 in endo-siRNA biogenesis.

## DATA AVAILABILITY

The atomic coordinates and structure factors for the DRH-3 N-terminal CARDs have been deposited with the Protein Data bank under accession code 6M6Q. The atomic coordinates and structure factors for the DRH-3 CTD domains have been deposited with the Protein Data bank under accession codes 6M6R and 6M6S.

## ACKNOWLEDGEMENT

This research is supported by Singapore Ministry of Health’s National Medical Research Council under its Open Fund – Individual Research Grant (OF-IRG) (NMRC/OFIRG/0075/2018.), the Singapore Ministry of Education under its Education Academic Research Fund Tier 1 (2018-T1-002-010) and Howard Hughes Medical Institute. We gratefully acknowledge the beamline staff at TPS 05A beamline in National Synchrotron Radiation Research Center, Hsinchu, Taiwan, MXII beamline in Australian Light Source, Melbourne, Australia and PSIII beamline in Swiss Light Source (SLS) Paul Scherrer Institut, Switzerland and NECAT 24-ID at Argonne National Laboratory, Argonne, IL, US for providing us outstanding support during the data collection. The authors wish to thank Drs. El Sahili Abbas and Chong Wai Liew for their help on X-ray data collection at synchrotron X-ray beamlines. The acknowledgements extend to the Protein Production Platform (PPP) at Nanyang Technological University School of Biological Sciences for assistance with high-throughput screening of protein overexpression. O.F. is a research scientist supported by the Howard Hughes Medical Institute. D.L. was a research fellow supported by the Howard Hughes Medical Institute. A.M.P. is an Investigator in the Howard Hughes Medical Institute.

