## Supplementary File for "Structural insights into the role of Dicer-related helicase 3 in RNAi in *Caenorhabditis elegans*"

### Supplementary Data

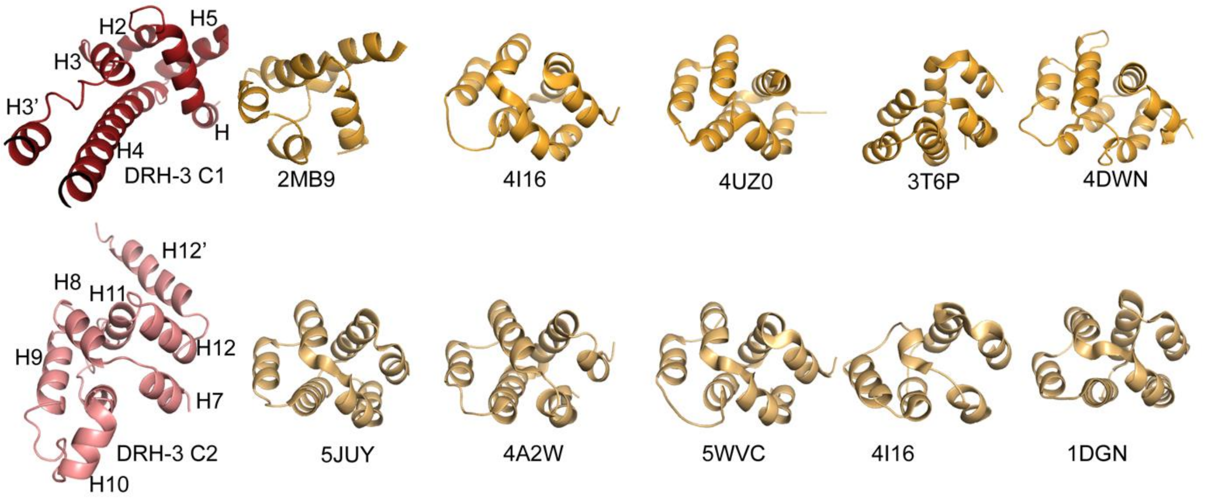


**Figure S1. Comparison of CARDs of DRH-3 NTD with top 5 structural hits from DALI search.** DRH-3 CARD1 and CARD2 are highlighted in red and salmon colours, respectively. The aligned regions of the 5 hits for each CARD are shown in orange colour. The six helices are labelled from N' terminus to C' terminus as H1-H6. The structures are listed, from *left* to *right*, in descending order of structure similarity by Z score, with the PDB code under the structure. Structural superimposition was done by pairwise DALI (1) and represented by PyMOL (2). For clarity, the structure models are rotated to give a similar orientation.

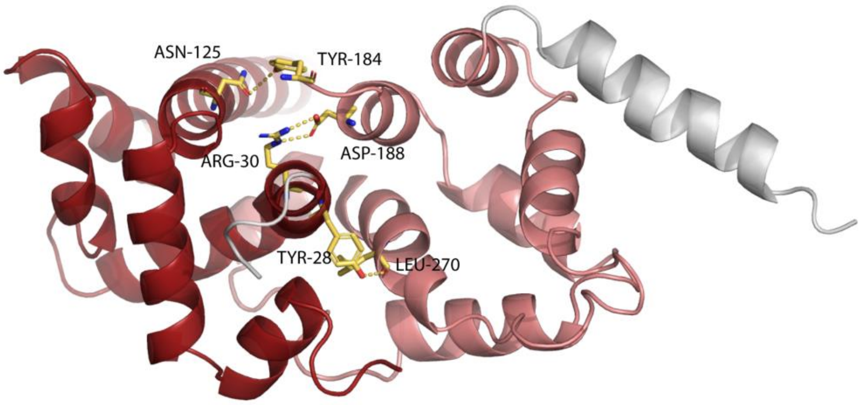


**Figure S2. The interface between the CARD1 and CARD2 of DRH-3.** The two CARDs in DRH-3 are arranged in parallel arrangement and the tandem CARDs are stabilized by hydrogen bonds. A close view of the interface is shown as cartoons. The interface residues are shown as sticks. The yellow dotted lines represent atomic distances of the polar bonds.

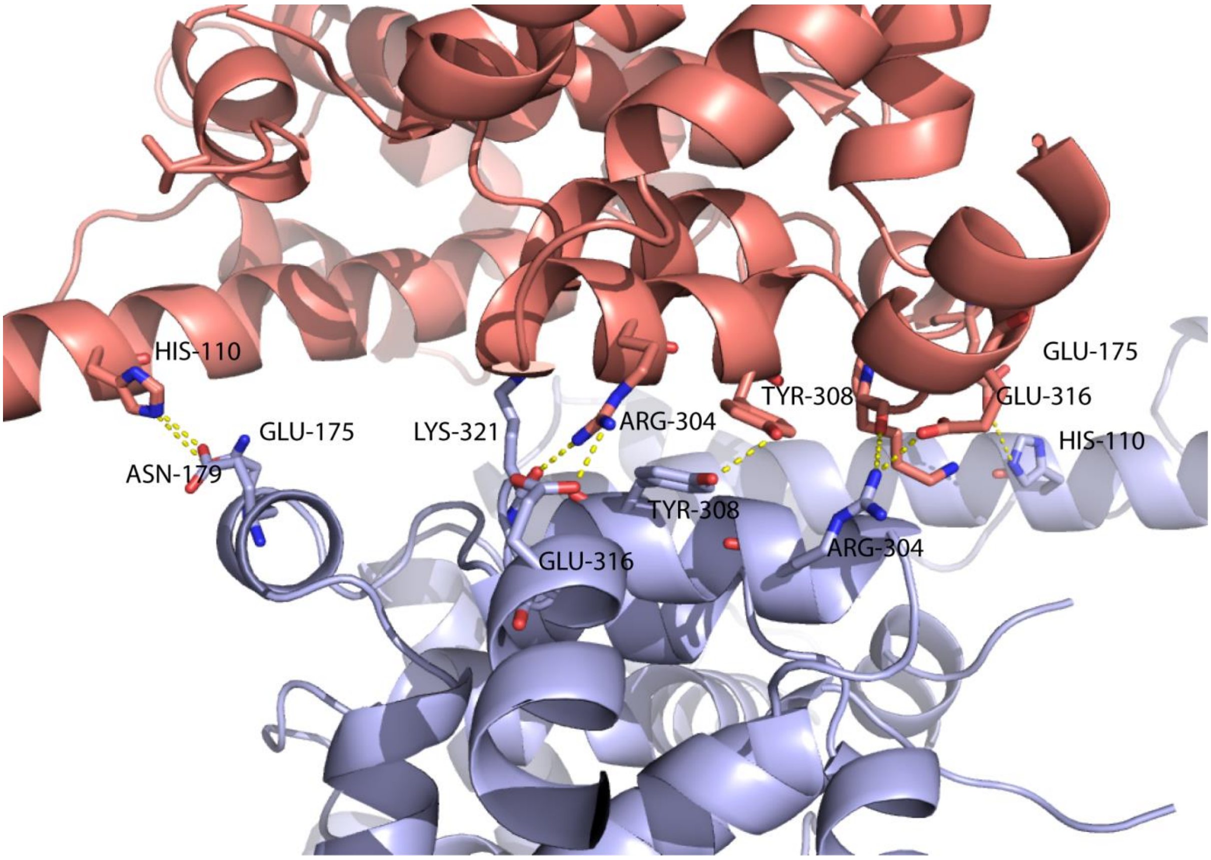


**Figure S3. Interactions involved in DRH-3 NTD dimer interface in the crystals.** We evaluated DRH-3 NTD for its ability to form dimer by PISA server. Four hydrogen bonds and six electrostatic interactions are involved in the dimer formation and the details are listed. A close-up view of the dimeric interface is shown as cartoons. The residues involved in dimerization are shown as sticks. The yellow dotted lines represent atomic distances of the polar bonds.

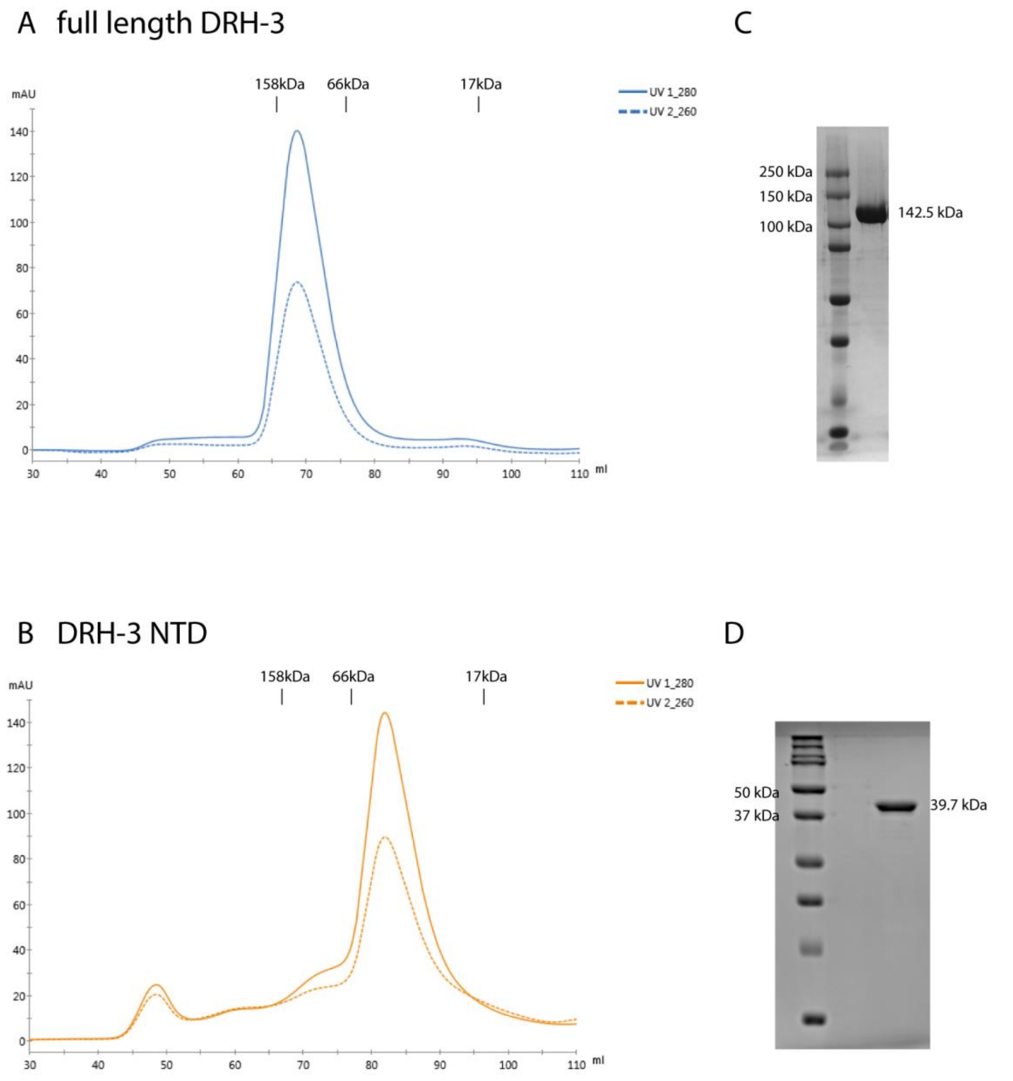


**Figure S4. Size exclusion chromatography profiles for full length DRH-3 (A) and DRH-3 NTD (B).** The Superdex 200 16/60 column (GE) was used for the characterization. The elution peaks indicate that both DRH-3 NTD and full length DRH-3 are in monomeric form in solution. Coomassie blue stained SDS-PAGE gel shows the purity of full length DRH-3 (C) and DRH-3 NTD (D).

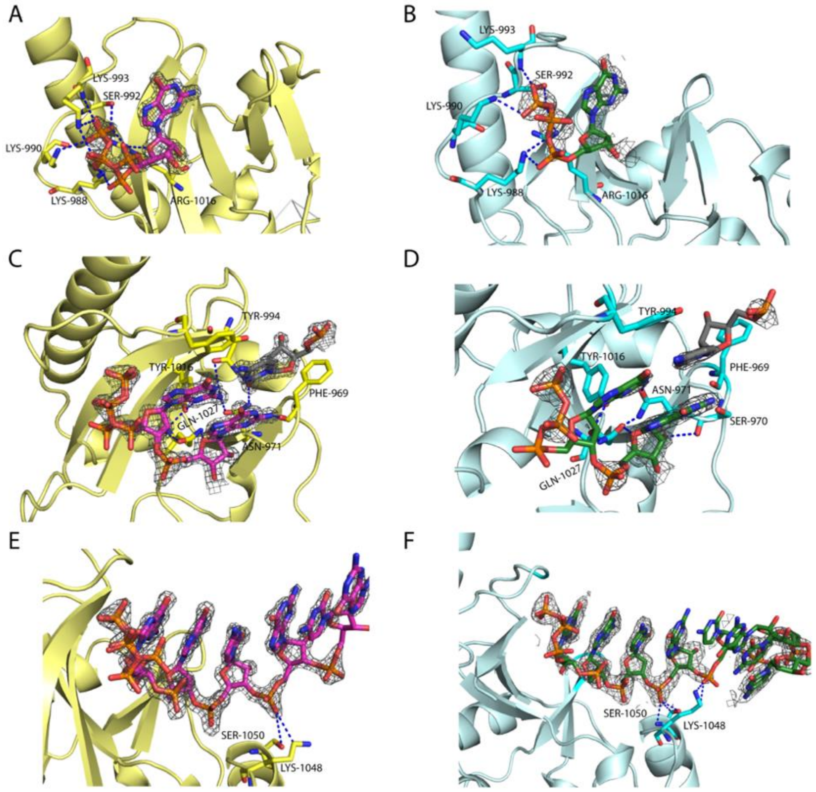


**Figure S5. Molecular details of DRH-3 CTD-RNA interactions.** (A) Interactions between the 5′-ppp of 12-mer dsRNA and DRH-3 CTD. The 5′-ppp12-mer dsRNA and DRH-3 CTD are depicted in magenta and yellow, respectively. (B) Interactions between the 5′-ppp of 8-mer ssRNA and DRH-3 CTD. The 5′-ppp 8-mer ssRNA and DRH-3 CTD are depicted in green and cyan, respectively. The 2Fo-Fc electron density map (gray mesh) for the 5'-ppp $G_1$ of the RNAs were contoured at 1.0 σ. The first nucleotide 5'-ppp $G_1$ of 5' strand and the key residues involved in RNA binding are shown as sticks. The dotted lines represent atomic distances of the polar bonds. K988, K990, S922, K933 and R1016 are the residues that recognize the 5'-ppp cap. (C,D) Interactions between the 5' $G_1G_2$ of the RNA and DRH-3 CTD. The electron density maps (2Fo-Fc omit maps contoured at 1.0 σ ) of $G_1$ and $G_2$ of the two RNAs were shown, respectively. The first two nucleotides $G_1$ and $G_2$ of 5' strand, the last nucleotides $C_8$ or $C_{12}$ of the 3' strand and the key residues involved in RNA binding are shown as sticks. The dotted lines represent atomic distances of the polar bonds. 5'-G1 is recognised mainly by Y1018, Q1027 and N971. N971 also recognizes 3'-C1 in dsRNA. Y994 and F969 contribute to the dsRNA stabilization by providing two hydrophobic interactions. (E,F) Interactions between the phosphate backbone of the RNA and the KWK motif of DRH-3 CTD. The electron density maps (2Fo-Fc omit maps contoured at 1.0 σ ) of the two RNAs were shown, respectively. The 5' strand RNAs and the key residues involved in RNA binding are shown as sticks. The dotted lines represent atomic distances of the polar bonds. In (DRH-3:5'-ppp 12-bp dsRNA)$^2$, C4 phosphate interacts with both K1048 and S1050. In DRH-3:5'-ppp 8-mer ssRNA, C4 phosphate makes contacts with S1050 while C5 phosphate forms interactions with LYS1048.

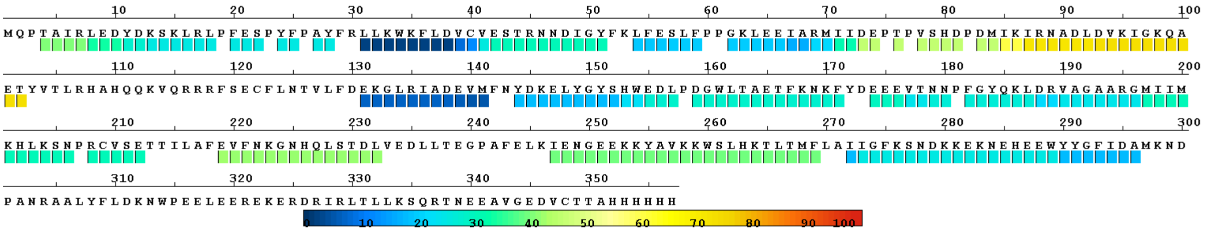


**Figure S6.** (Related to Figure 4.) **HDX-MS heat map of DRH-3 NTD.** Percentage change in HDX uptake (scale at the bottom) is plotted on peptides analysed in the experiment.

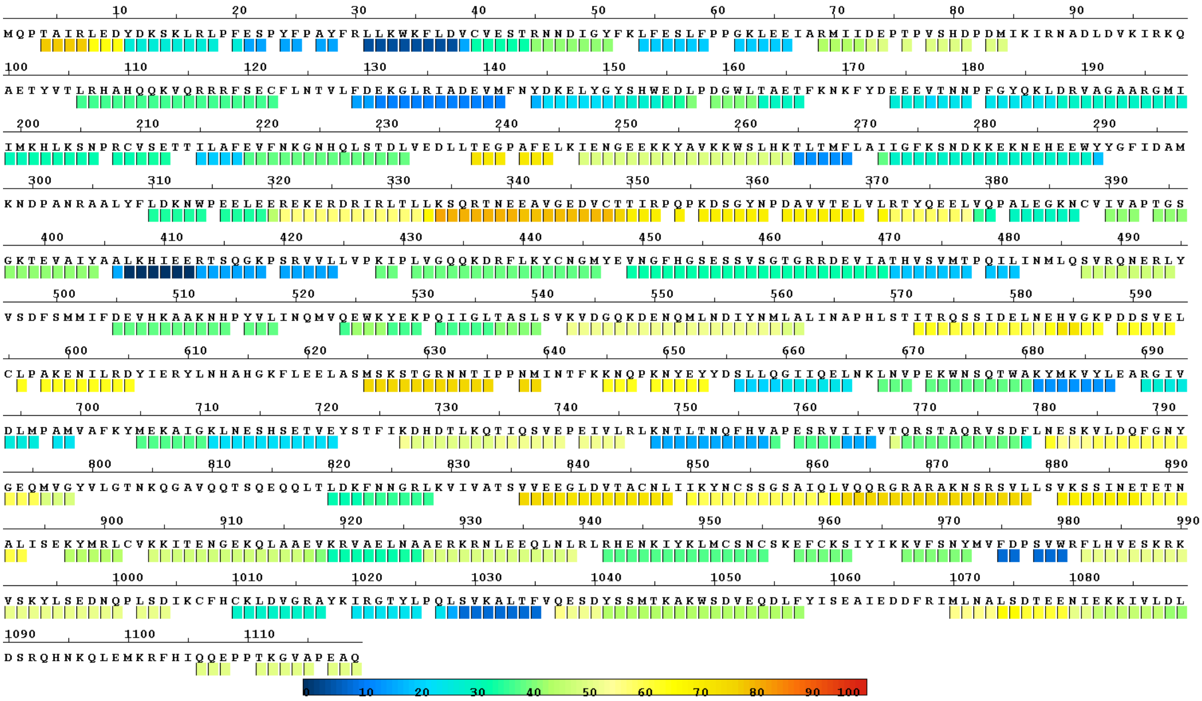

**Figure S7.** (Related to Figure 4.) **HDX-MS heatmap of full length DRH-3.** Percentage change in HDX uptake (scale at the bottom) is plotted on peptides analysed in the experiment.

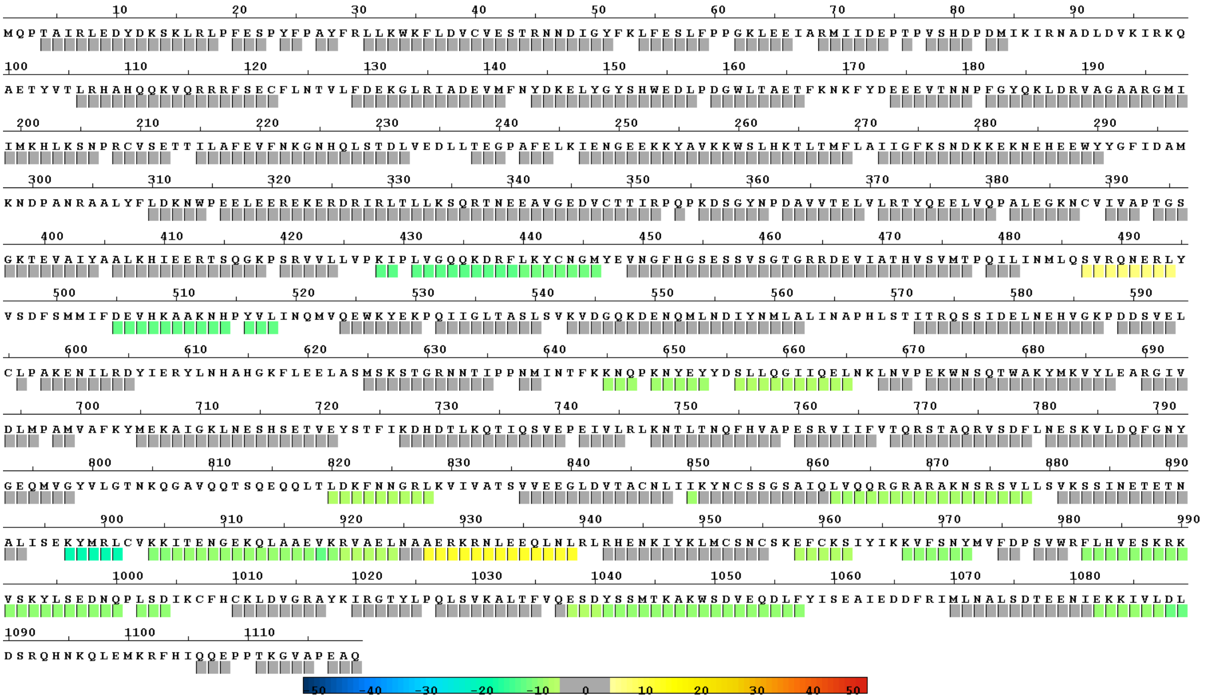

**Figure S8.** (Related to Figure 4.) **HDX-MS difference map of full length DRH-3 binding to 3P10L.** Percentage change in HDX uptake (scale at the bottom) is plotted on peptides analysed in the experiment.

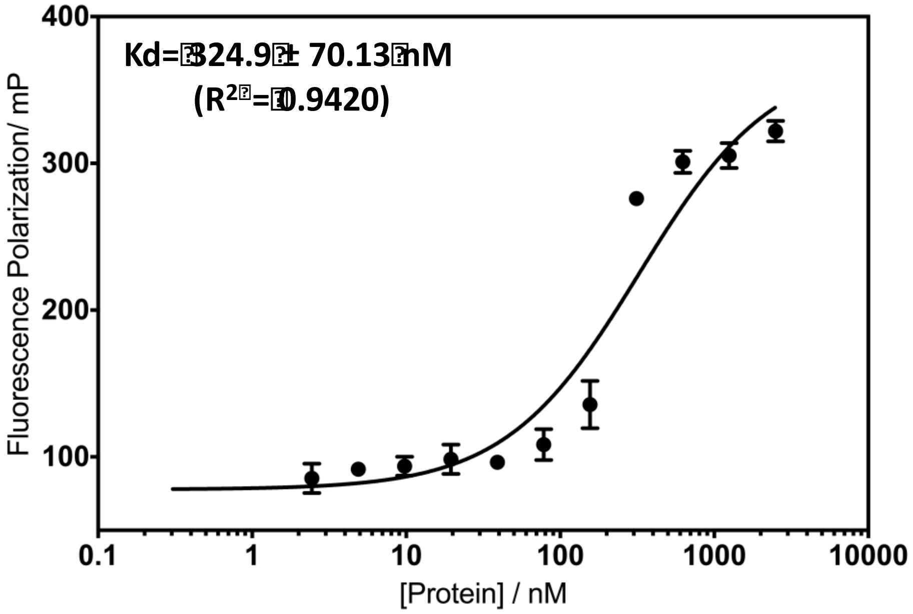


**Figure S9.** Direct FP binding assay to determine $Kd$ of the DRH-3 CTD. To characterize the baseline FP signal, the proteins were serially diluted in the concentration range of 1–5000 nM. The FP signals were measured and plotted as a function of protein concentration. Average values ± standard error (SE) from three measurements per condition are shown in the graph.

**Table S1. Top five DALI hits for DRH-3 CARDs.**

| PDB code | chain | Z | R.m.s.d | Length aligned | No. of residues | Identity (%) | Description |
|---|---|---|---|---|---|---|---|
| TOP 5 HITS FOR DRH-3CARD1 | | | | | | | |
| PDB code | chain | Z | R.m.s.d | Length aligned | No. of residues | Identity (%) | Description |
| 2mb9 | A | 5.5 | 3.2 | 92 | 106 | 10 | B-CELL LYMPHOMA/ LEUKEMIA 10 |
| 4i16 | A | 4.9 | 3.2 | 81 | 90 | 11 | CASPASE RECRUITMENT TO DOMAIN-CONTAINING PROTEIN 11 |
| 4uz0 | A | 4.9 | 3.7 | 80 | 87 | 8 | NUCLEOLAR PROTEIN 3 |
| 3t6p | A | 3.7 | 3 | 78 | 330 | 13 | BACULOVIRAL IAP REPEAT-CONTAINING PROTEIN 2 |
| 4dwn | A | 3.6 | 3.2 | 80 | 97 | 8 | BCL10-INTERACTING CARD PROTEIN |
| TOP 5 HITS FOR DRH-3 CARD2 | | | | | | | |
| PDB code | chain | Z | R.m.s.d | Length aligned | No. of residues | Identity (%) | Description |
| 5juy | B | 6.4 | 6.1 | 69 | 1234 | 15 | APOPTOTIC PROTEASE-ACTIVATING FACTOR 1 |
| 4a2w | A | 5.8 | 3.2 | 85 | 678 | 11 | RETINOIC ACID INDUCIBLE PROTEIN I |
| 5wcv | B | 5.5 | 2.9 | 87 | 103 | 10 | APOPTOTIC PROTEASE-ACTIVATING FACTOR 1 |
| 4i16 | A | 5.4 | 3.0 | 87 | 90 | 8 | CASPASE RECRUITMENT TO DOMAIN-CONTAINING PROTEIN 11 |
| 1dgn | A | 5.0 | 2.8 | 80 | 89 | 9 | ICEBERG (PROTEASE INHIBITOR) |

#### Table S2. List of RNAs.

**Chemical synthesis**

| **Name** | RNA sequence |
|---|---|
| **R0** | U-FAM<br>5′OH-UUUAUAUAUUAGGACCGC \\<br>AAAUAUAUAAUCCUGGCG C<br>G |
| **R1** | 5′ppp-GGCGCGCGCC<br>CCGCGCGCGG-5′ppp |
| **R1OH** | 5′OH-GGCGCGCGCC<br>CCGCGCGCGG-5′OH |
| **R1P** | 5′p-GGCGCGCGCC<br>CCGCGCGCGG-5′ppp |
| **R2** | 5′pppGGGCGCGCGCC<br>CCGCGCGCGGG-5′ppp |
| **R3** | 5′pppGGCGCGCGCCC<br>CCCGCGCGCGG-5′ppp |

**In vitro transcription**

| **Name** | RNA sequence | DNA sequence |
|---|---|---|
| **3P10L** | U<br>5′PPP-GGACGCGUGC U<br>CCUGCGCACG C<br>G | **5′-GTAATACGACTCACTATA**GGAGAGGUGCUUCGGCACGCGUCC |
| **8-nt ssRNA** | 5'ppp-GGCCGCCC | **5′-GTAATACGACTCACTATA**GGCCGCCC |
| **12-mer dsRNA** | 5′ppp-GGCGCGCGCC<br>CCGCGCGCGG-5′ppp | **5′-GTAATACGACTCACTATA**GGCGCGCGCC |
| **21-mer hairpin RNA with 3'-UU** | 5′ppp-UGAGGUAGUAGGUUGUAUAGUUUGCAAACUAUACAACCUACUACCUCAUU | **5′-**<br>**GTAATACGACTCACTATA**TGAGGTAGTAGGTTGTATAGTTTGCAAACTATACAACCTACTACCTCATT |
| **100-mer dsRNA** | **5′PPP-**<br>**GGAGAAAAGCUGUUUGUCUCCCCAGCAUACUUUAUCGCCUUCACUGCCUUUUGACUGCAGAGGAGC**<br>**UAAGAGCCCAGACUGCUAUAGUGAGUCGUAUUAC-3′**<br><br>3′-<br>CCUCUUUUCGACAAACAGAGGGGUCGUAUGAAAUAGCGGAAGUGACGGAAAACUGACGUCUCCUCG<br>AUUCUCGGGUCUGACGAUAUCACUCAGCAUAAUG-5′PPP | **5′-**<br>**GTAATACGACTCACTATA**GGAGAAAAGCTGTTTGTCTCCCCAGCATACTTTATCGCCTTCACTGCCTTTT<br>GACTGCAGAGGAGCTAAGAGCCCAGACTGCTATAGTGAGTCGTATTAC<br>3′-<br>CATTATGCTGAGTGATATCCTCTTTTCGACAAACAGAGGGGTCGTATGAAATAGCGGAAGTGACGGAAAA<br>CTGACGTCTCCTCGATTCTCGGGTCTGACGA**TATCACTCAGCATAATG** |
